# Generation of appropriate protein structures for virtual screening using AlphaFold3 predicted protein–ligand complexes

**DOI:** 10.1101/2025.02.17.638750

**Authors:** Yuki Yasumitsu, Masahito Ohue

**Affiliations:** Department of Computer Science, School of Computing, Institute of Science Tokyo, 4259 Nagatsuta-cho, G3-56 Midori-ku, Yokohama, 226-8501, Kanagawa, Japan

**Keywords:** AlphaFold3, Virtual Screening, Structure-Based Virtual Screening, Protein–Ligand Docking, Protein Structure Prediction

## Abstract

In early drug discovery, virtual screening—a computational method for selecting candidate compounds—helps reduce development costs. Traditionally, structure-based virtual screening required experimental protein structures, but advances like AlphaFold2 have begun to overcome this limitation. However, AlphaFold2 does not capture ligand-induced conformational changes (transition from apo to holo forms), limiting its utility for protein–ligand docking. In this study, we evaluate AlphaFold3, which predicts protein–ligand complex structures when both protein and ligand inputs are provided. Using the DUD-E dataset and Uni-Dock, we show that holo structures predicted with ligand inclusion yield higher screening performance than apo structures generated without ligand input. Notably, incorporating active ligands enhances screening performance, whereas inactive (decoy) ligands produce results similar to apo predictions. The use of template structures further improves outcomes. We also analyze the impact of ligand molecular weight, binding pocket location, and AlphaFold3 ranking scores on screening performance. Our findings indicate that lower molecular weight ligands tend to generate predicted structures that more closely resemble experimental holo structures, thus improving screening efficacy. Conversely, larger ligands (700–800) can induce open binding pockets that favor screening for some targets. These results suggest that employing AlphaFold3 with appropriate ligand inputs is a promising strategy for virtual screening, particularly for proteins lacking experimental structural data.

## 1. Introduction

Virtual screening (VS) is a computational method used to select candidate drug compounds from a vast chemical space [1, 2](estimated to be between 10^30^ and 10^60^ compounds [3]). In drug discovery, such computational filtering is essential for reducing costs. VS methods can be broadly categorized into ligand-based [4] and structure-based [5] approaches, with the latter evaluating candidate compound activity based on the three-dimensional structure of the target protein. In structure-based virtual screening (SBVS), compounds are ranked by predicting their binding affinity, and protein–ligand docking simulations [6] or molecular dynamics simulations [7] are used as methods for predicting this binding affinity.

Unlike ligand-based methods, which rely on experimental data from known active compounds, SBVS does not require prior experimental information on active compounds, making it well-suited for discovering novel drug candidates. However, protein structures often change conformation upon ligand binding. Structures without a bound ligand (apo structures) generally perform worse in docking simulations than structures with a bound ligand (holo structures) [8, 9].

When the experimental structure of a target protein is unavailable, the protein’s structure must be predicted. Although homology modeling has been used traditionally, its dependence on suitable template structures remains a limitation [10, 11, 12]. Recent advances in machine learning-based protein structure prediction—notably AlphaFold2 [13]—have shown promise for SBVS. However, studies have reported that AlphaFold2 predicted structures yield virtual screening performance comparable to apo structures rather than holo structures [9, 14, 15].

Released in 2024, AlphaFold3 [16] is capable of predicting protein–ligand complexes directly. It is hypothesized that the holo structures predicted by AlphaFold3 will exhibit improved screening performance compared to those predicted by AlphaFold2. Yet, the evaluation of AlphaFold3 for virtual screening and the determination of optimal ligand input remain open questions. In this study, we evaluate the screening performance of AlphaFold3-predicted holo structures and investigate ligand input selection strategies to obtain structures with enhanced screening performance.

## 2. Materials and Methods

### 2.1. AlphaFold3 Prediction of Holo Structures

To evaluate the screening performance of holo structures predicted by AlphaFold3, we compared the performance of different input conditions. Figure 1A shows a schematic diagram of holo structure prediction by AlphaFold3. We generated:

**Figure 1.**
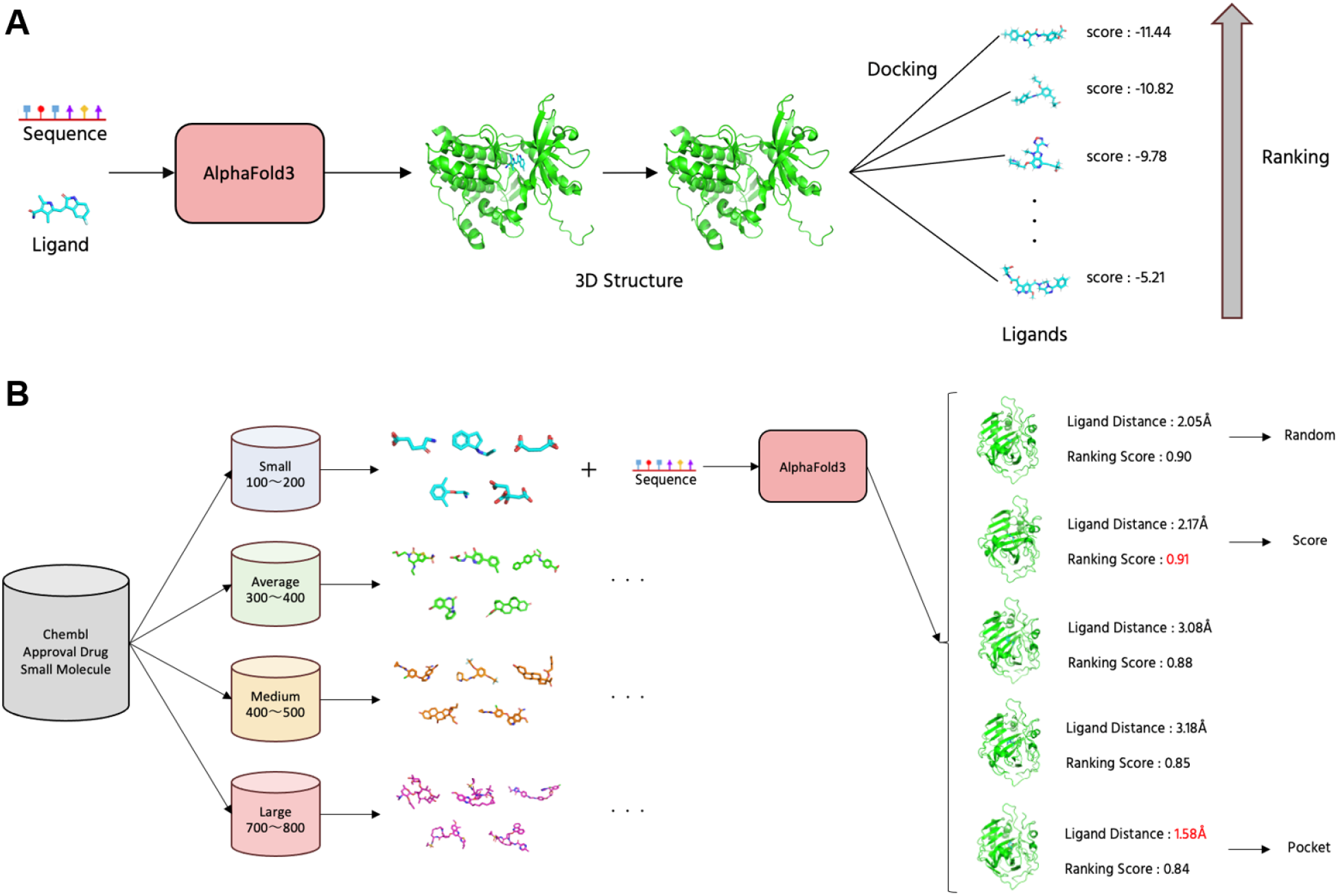
Overview of the proposed method. (A) Schematic diagram of holo structure prediction using AlphaFold3. (B) Schematic diagram of the ligand input selection strategy for AlphaFold3.

- **apo**: An apo structure predicted by providing only the protein sequence (i.e., without any ligand),
- **holo crystal**: A holo structure predicted using the ligand from the corresponding PDB crystal structure. Since this ligand is part of AlphaFold3’s training data, a high prediction accuracy is expected,
- **holo active**: A holo structure predicted by randomly selecting an active ligand from the DUD-E dataset,
- **holo decoy**: A holo structure predicted by randomly selecting an inactive (decoy) ligand from DUD-E.

Furthermore, for each of these four types, predictions were performed both with and without the use of default template structures (denoted as *tmpl*), resulting in a total of eight prediction types.

### 2.2. Ligand Input Selection Strategy in AlphaFold3

Since the predicted holo structure and its subsequent screening performance may vary depending on the ligand used as input, we investigated different ligand input selection strategies. Figure 1B illustrates the overall strategy. As the ligand dataset, we used approved small-molecule drugs from ChEMBL [17]. Out of 2477 compounds, 2336 compounds with successfully computed molecular weights (using RDKit) were used (average molecular weight: 363.7; median: 337.6).

We grouped the ligands based on molecular weight as follows:

- **small**: 100–200,
- **average**: 300–400,
- **medium**: 400–500,
- **large**: 700–800.

For each group and target, three types of holo structures were predicted:

- **random**: A holo structure generated using one randomly selected ligand,
- **pocket**: Among five randomly selected ligands, the holo structure whose binding pocket position is closest to that of the crystal structure is chosen. This is evaluated by computing the Ligand Distance (i.e., the distance between the centers of the ligand in the predicted and crystal structures),
- **score**: Among five randomly selected ligands, the holo structure with the best AlphaFold3 ranking score is selected.

### 2.3. Virtual Screening

The DUD-E dataset [18] comprises targets with associated active and decoy compounds and is widely used as a benchmark for virtual screening. In this study, we used 87 targets consisting of single-chain pocket from the DUD-E dataset (excluding 4 targets where AlphaFold3 predictions failed). Docking simulations were performed using Uni-Dock [19], a GPU-accelerated protein–ligand docking tool that is computationally efficient for large-scale virtual screening. The predicted structure was aligned with the crystal structure using PyMOL[20]. Additionally, AutoDockTools[21] was used for the protein preprocessing.

### 2.4. Evaluation Metrics

#### 2.4.1. Structural Similarity

We evaluated the structural similarity of the predicted models using the Root Mean Square Deviation (RMSD), computed for both the entire structure and the binding pocket region:

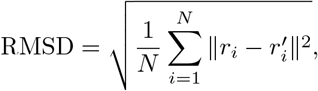

where *ri* and 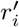 represent the coordinates of the *i*th atom in the predicted and reference structures, respectively.

#### 2.4.2. Binding Pocket Location

The binding pocket position was evaluated by computing the Ligand Distance, defined as the distance between the center coordinates of the ligand in the predicted structure and those in the crystal structure after alignment.

#### 2.4.3. Screening Performance

Screening performance was assessed using the Receiver Operating Characteristic Area Under the Curve (ROC-AUC) and the Enrichment Factor at 1% (EF1%). ROC-AUC measures the predictive accuracy of a binary classifier (with values closer to 1 indicating better performance). The EF quantifies the enrichment of active compounds in the top-ranked portion of the screening results:

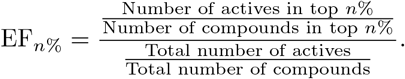

### 2.5. Experimental Environment

The following software versions were used:

- PyMOL: 2.5.5,
- Uni-Dock: 1.1.1,
- AutoDockTools: 1.5.7,
- RDKit: 2024.03.6.

## 3. Results

### 3.1. AlphaFold3 Holo Structure Prediction and Screening Results

For the DUD-E dataset, we generated eight types of predicted structures (apo, holo crystal, holo active, holo decoy, with and without templates) using AlphaFold3 and evaluated their screening performance. Figure 2A shows the ROC-AUC results, while Figure 2B presents the EF1% results. The performance ranking was as follows: holo crystal yielded the best screening performance, followed by holo active, holo decoy, and apo. Although the holo crystal predictions achieved screening performance comparable to the holo PDB structures, both holo active and holo decoy exhibited lower performance than the holo PDB. In particular, holo decoy showed performance only marginally better than or comparable to apo. In addition, using template structures improved the screening performance. Figures 2C and 2D show the overall RMSD and pocket RMSD between the DUD-E holo PDB structures and the predicted structures, respectively, indicating that holo crystal is most similar to the experimental structure, with holo active following.

**Figure 2.**
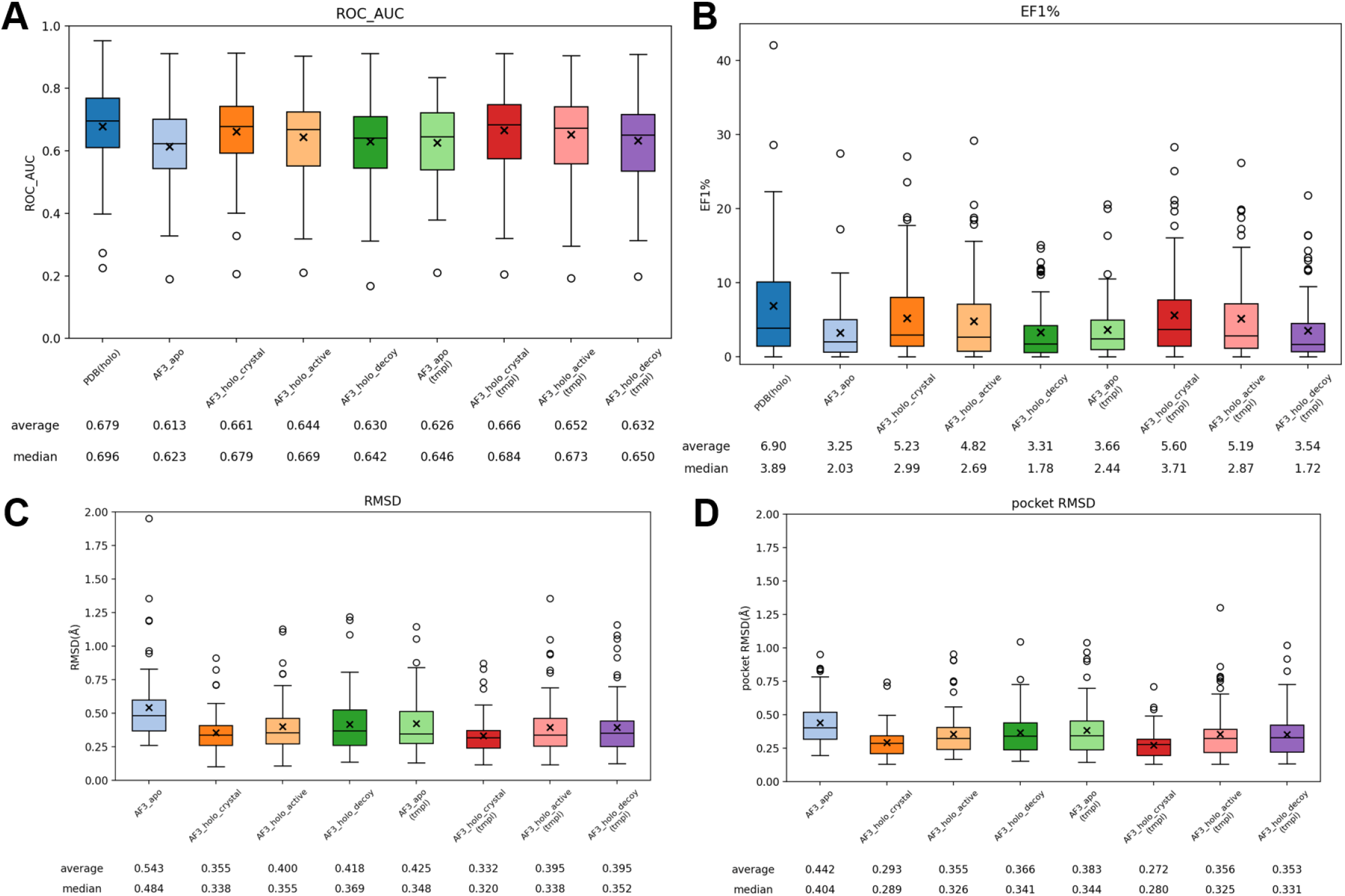
Virtual screening performance and structural similarity of predicted structures by AlphaFold3. (A) ROC-AUC evaluation, (B) EF1% evaluation, (C) Overall RMSD between DUD-E holo PDB structures and predictions, (D) Pocket RMSD between DUD-E holo PDB structures and predictions.

### 3.2. Ligand Input Selection Strategy in AlphaFold3

For the DUD-E dataset, we generated 12 sets of predictions by combining four ligand size groups (small, average, medium, large) with three selection methods (random, pocket, score) without using templates. Figure 3A and Figure 3B show the virtual screening performance evaluated by ROC-AUC and EF1%, respectively. Figure 3C displays the pocket RMSD, and Figure 3D shows the Ligand Distance. The ROC-AUC was highest for the medium group, followed by large and average, with the small group being the lowest. The EF1% values increased in the order of small, average, and medium, while the large group was slightly higher than medium and comparable to average. Furthermore, both the pocket and score selection methods yielded better performance than random selection. The pocket RMSD was similar for small and average, slightly worse for medium, and worst for large ligands, indicating that larger ligands tend to produce predicted structures that deviate from the PDB holo structures. In terms of Ligand Distance, the average and medium groups performed best, while the small group performed poorly and the large group was second worst, suggesting that ligands with molecular weights between 300 and 500 yield higher pocket positioning accuracy.

**Figure 3.**
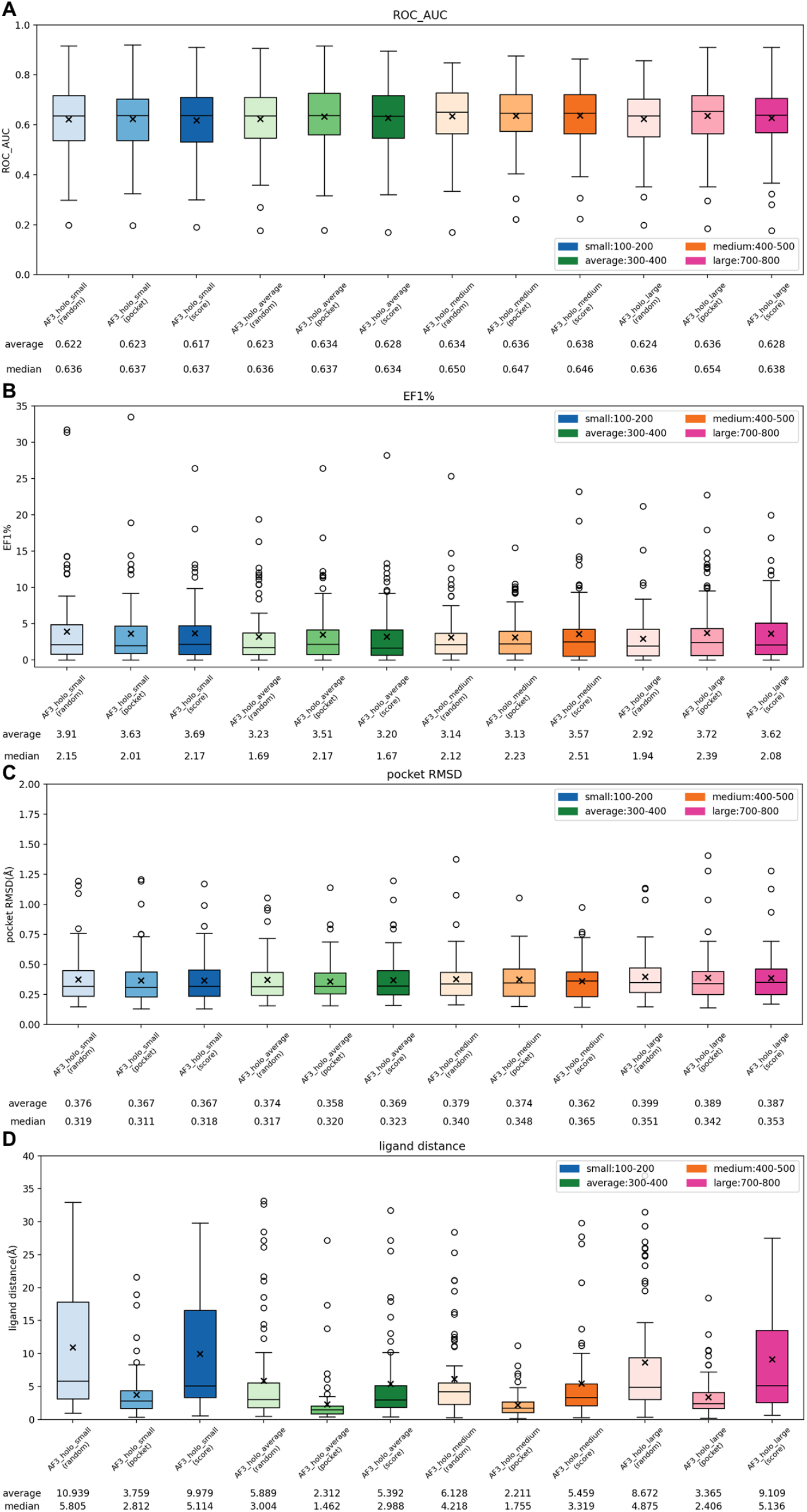
Virtual screening performance and structural similarity when varying the input ligand size in AlphaFold3. (A) ROC-AUC evaluation, (B) EF1% evaluation, (C) Pocket RMSD between DUD-E holo PDB structures and predictions, (D) Ligand Distance between DUD-E holo PDB structures and predictions.

### 3.3. Predicted Structures and ROC Curves

Figure 4 presents the ROC curves and structure overlays for representative targets. In Figure 4A, the overlay of the crystal structure and the predicted structure for THB, along with its ROC curve, demonstrates that using an active ligand results in a predicted structure closer to the holo PDB structure and improved screening performance compared to the apo structure. Figures 4B and 4C show examples for RXRA and KIF11, respectively. For both targets, larger ligand inputs led to an open binding pocket conformation; while RXRA benefited in terms of screening performance, KIF11 exhibited a decrease. Thus, although large ligand inputs can generate an open binding pocket conformation, they do not necessarily guarantee improved virtual screening accuracy.

**Figure 4.**
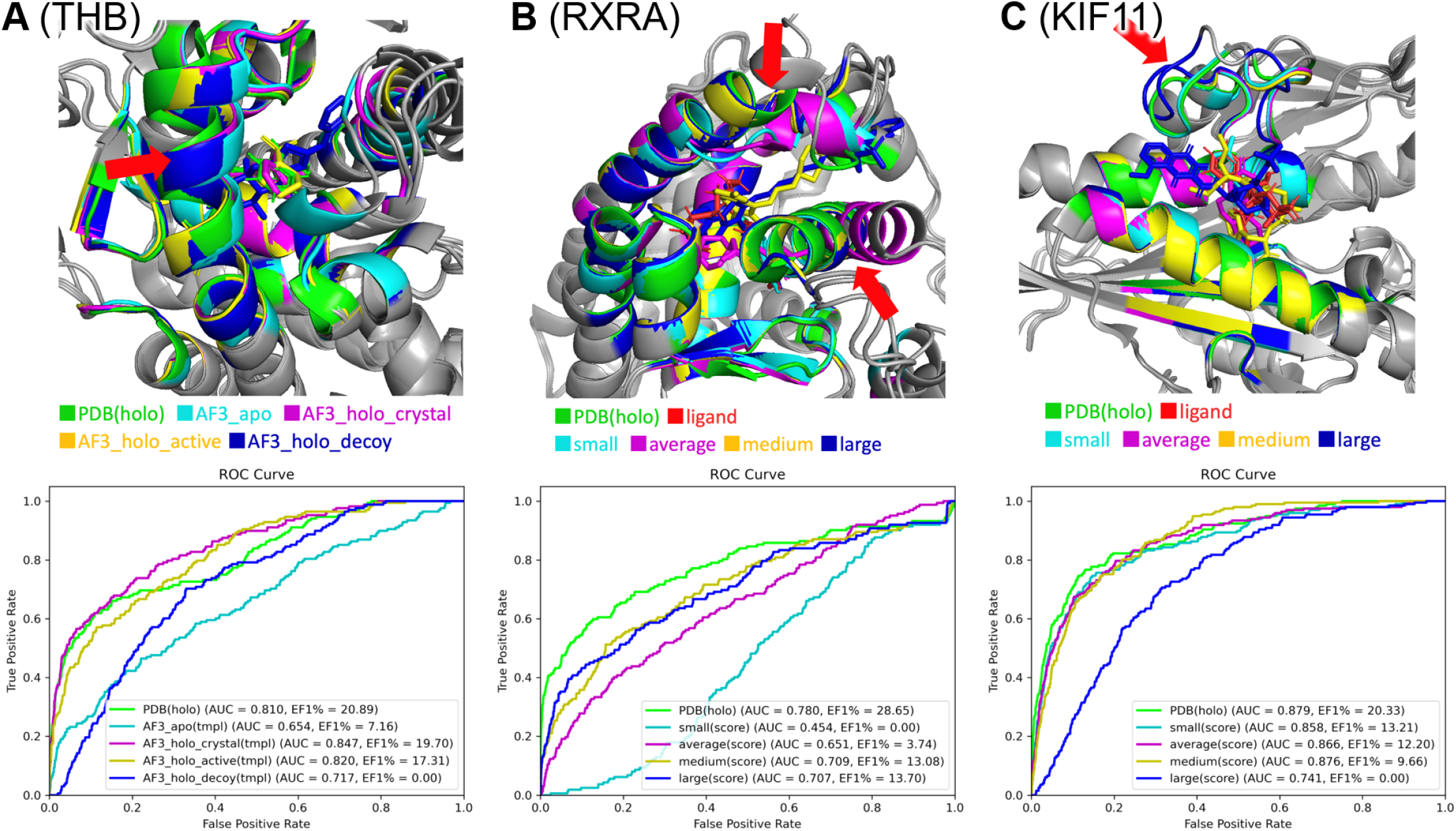
Overlay of predicted structures and ROC curves for representative targets. (A) THB, (B) RXRA, (C) KIF11.

## 4. Discussion

### 4.1. Evaluation of Screening Performance Using AlphaFold3 Holo Structures

We evaluated the prediction accuracy and screening performance of four types of structures (apo, holo crystal, holo active, holo decoy) predicted by AlphaFold3. The results demonstrated that holo crystal yielded the best screening performance, followed by holo active, holo decoy, and apo. This confirms that predicting holo structures improves screening performance. Moreover, the choice of input ligand affects both the predicted holo structure and the subsequent screening performance. While the ligand from the crystal structure (which is part of the training data) yields very high prediction accuracy, it is less applicable for model evaluation. Comparing holo active and holo decoy, the active ligand produced better prediction accuracy and screening performance, suggesting that using an active ligand is beneficial. However, holo decoy only achieved marginal improvements or performance comparable to apo, indicating a challenge when the active ligand is unknown.

### 4.2. Optimal Ligand Input Selection

Among the three ligand selection methods (random, pocket, score), both the pocket and score methods – which select the best structure from five randomly chosen ligands based on pocket position or ranking score – outperformed random selection. In practice, since the binding pocket position may not be known a priori, it is desirable either to combine this approach with a pocket prediction tool or to base the selection on the ranking score. Furthermore, the optimal ligand molecular weight appears to be target-dependent. Thus, strategies such as ensemble docking using multiple predicted structures or adapting the input ligand based on the docking ligand’s size may be required.

## 5. Conclusion

Using AlphaFold3, we predicted holo structures and evaluated their prediction accuracy and virtual screening performance. The results show that holo predictions exhibit higher screening performance compared to apo predictions. In addition, using an active ligand as input improves screening performance, whereas inactive ligands yield performance comparable to apo structures. By varying the ligand molecular weight, we found that smaller ligands tend to produce predicted structures closer to the PDB holo structure and yield higher screening performance. Conversely, larger ligands induce an open pocket conformation that improves screening performance for some targets. Moreover, considering the pocket location and the AlphaFold3 ranking score further enhances screening outcomes. Although our evaluation was performed using the DUD-E dataset (which is included in AlphaFold3’s training data), further evaluation on targets outside the training set is warranted.

## Declaration of Generative AI and AI-assisted technologies in the writing process

No declarations were made.

## Author contributions

Yuki Yasumitsu: Conceptualization, Methodology, Software, Validation, Formal analysis, Investigation, Resources, Data curation, Visualization, Writing – original draft, Writing – review & editing. Masahito Ohue: Conceptualization, Methodology, Formal analysis, Investigation, Writing – review & editing, Supervision, Project administration, Funding acquisition.

## Acknowledgment

Computational experiments were performed using the TSUBAME 4.0 supercomputer at the Institute of Science Tokyo.

